# Cyanobacteria newly isolated from marine volcanic seeps display rapid sinking and robust, high density growth

**DOI:** 10.1101/2023.10.30.564770

**Authors:** Max G. Schubert, Tzu-Chieh Tang, Isabella M. Goodchild-Michelman, Krista A Ryon, James R. Henriksen, Theodore Chavkin, Yanqi Wu, Teemu P. Miettinen, Stefanie Van Wychen, Lukas R. Dahlin, Davide Spatafora, Gabriele Turco, Michael T. Guarnieri, Scott R. Manalis, John Kowitz, Raja Dhir, Paola Quatrini, Christopher E Mason, George M Church, Marco Milazzo, Braden T Tierney

## Abstract

Cyanobacteria are photosynthetic organisms that play important roles in carbon cycling as well as promising bioproduction chassis. Here, we isolate two novel cyanobacteria, UTEX 3221 and UTEX 3222, from a unique marine environment with naturally elevated CO₂. We describe complete genome sequences for both isolates and, focusing on UTEX 3222 due to its planktonic growth in liquid, characterize biotechnologically-relevant growth and biomass characteristics. UTEX 3222 outpaces other fast-growing model strains on solid medium. It can double every 2.35 hours in a liquid medium and grows to high density (>31g/L biomass dry weight) in batch culture, nearly double that of *Synechococcus* sp. PCC 11901, whose high-density growth was recently reported. In addition, UTEX 3222 sinks readily, settling more quickly than other fast-growing strains, suggesting improved de-watering of UTEX 3222 biomass. This settling behavior can be explained in part by larger cell volume. These traits may make UTEX 3222 a compelling choice for photosynthetic bioproduction from CO₂. Overall, we find that bio-prospecting in environments with naturally elevated CO₂ may uncover novel CO₂-metabolizing organisms with unique characteristics.

## Introduction

Cyanobacteria serve as promising hosts for photosynthetic bioproduction^1–4^. They carry out oxygenic photosynthesis, which is widely understood to be the most important metabolic innovation in Earth’s history^5^. Using CO₂ as a carbon source, light as an energy source, and water as an electron donor, they build complex living materials, converting ubiquitous materials to diverse substrates.

Cyanobacteria potentially enable a variety of new carbon-negative technologies^6–8^. Most cyanobacterial research is conducted in model organisms isolated more than 50 years ago^9,10^, but more recently-isolated strains from diverse environments demonstrate unique traits and potentially improved biotechnological potential. For example, UTEX 2973 is a spontaneous mutant of the historical type strain PCC 6301^11^, exhibiting high light tolerance and exceptional doubling times as fast as every 1.5 hours^12–14^. PCC 11801 and PCC 11802 were isolated from a eutrophic urban lake and display fast growth and promising metabolic traits^15–17^. PCC 11901, isolated from a fish farm, displays fast growth to unusually high biomass density^18,19^. Cyanobacteria isolated from alkaline soda lakes grow well at high pH, which is beneficial for CO_2_ transfer into water^20,21^. Isolation of novel cyanobacteria with potentially improved industrial character is thus a promising area of recent research.

We hypothesized that, due to their exposure to sunlight and high ambient dissolved inorganic carbon, the shallow volcanic seeps off the coast of Baia di Levante in Vulcano Island, Italy, may be rich in biotechnologically relevant cyanobacterial life. This volcanic region features marine volcanic seeps, which continuously release CO₂, and is actively investigated as a model of ocean acidification and ecosystem structure^22^. The Baia di Levante is well studied for its shallow seeps at 1-4 m depth, discharging ∼1300 tons/year^23^. For comparison, the few existing flux estimates for volcanic CO_2_ seeps worldwide range from <1 to > 2,000,000 tons/year^24^. The CO_2_-rich emissions at Baia di Levante result in acidic conditions (<6.5 pH) in the seawater column around the main venting area. The discharged fluids consist of hydrothermal gasses containing elevated CO_2_ (>98%), H_2_S (400 ppm), and CH_4_ (400 ppm) concentrations. The interaction of reduced gasses with seawater leads to dissolved oxygen consumption and reducing conditions (low redox potential, Eh) in seawater^25,26^. Such reducing conditions are also caused by the discharge into seawater of saline hydrothermal brines (derived from a shallow aquifer). These fluids are rich in metals such as iron, whose oxidation leads to extensive oxygen consumption^25,26^. Iron concentrations in the bay and nearby the main degassing area are roughly an order of magnitude higher than control waters^27^.

In contrast to the deeper oceanic vents that sunlight does not reach, the shallow reaches have water, light, and CO₂ — all crucial for oxygenic photosynthesis — in copious abundance. Carbon is often a limiting factor in cyanobacterial growth in the environment, and high-affinity cyanobacterial carbon concentration mechanisms have evolved to mitigate this limitation^28^. We hypothesized that organisms relieved from carbon limitation would potentially realize greater fitness improvement from, and thus evolve more readily, adaptations addressing other limitations. These could include efficient light utilization, rapid growth, and division, evasion of predators, antagonism toward competitors, or countless other possibilities. By isolating organisms from this unique environment, we can expect to discover unique organisms valuable for research and capable of sequestering carbon with high efficiency.

In this work, we endeavored to isolate such organisms, with a focus on cyanobacteria, to contribute to the growing set of novel, promising cyanobacterial hosts for bioproduction. Here, we describe the isolation, observation, and genome sequencing of two related strains found in this unique environment. We additionally describe the initial growth and biomass characterization of one of these strains, displaying substantial biotechnological potential. These new isolates are well-suited to rapid culture in the lab, growing quickly to high biomass density rivaling that of the unusually high density previously observed in PCC 11901. These strains also display larger cell sizes than other fast-growing cyanobacteria and sink more readily, suggesting potential improvements to industrial dewatering of biomass through centrifugation, settling, or filtration. These newly-isolated strains are a compelling new chassis to consider for photosynthetic production of foods, chemicals, or even as a route for carbon sequestration.

## Results

### Obtaining and Sequencing Fast-Growing Cyanobacteria

An expedition to multiple marine zones near Vulcano Island (Aeolian archipelago, Italy) was conducted due to the unique CO_2_-enriched environment present^22^. We selected eight sites in Baia di Levante where a pH/pCO_2_ gradient extends over 500m from a shallow main venting area (latitude longitude 38.418722N, 14.963879E), generating a seawater pH of approximately 6.5, surface temperatures of approximately 25°C and pCO_2_ averaging between 14,500 µatm and 28,800 µatm in its surrounding waters (Table S1).

Sediment, water, and biomass samples for microbiological analysis were obtained on open-circuit SCUBA from these zones (Fig. 1a, *Methods*). This study aimed to isolate fast-growing cyanobacteria from such samples. Multiple seawater and sediment samples were pooled from each dive site, concentrated by filtration, enriched in conditions expected to promote fast liquid growth of phototrophs, and rendered axenic on solid medium (*Methods*). Two promising cyanobacterial isolates were obtained from two sites from the shallow CO2 seep area of Baia di Levante and are now publicly available as UTEX 3221 and UTEX 3222. Both strains grew well on solid medium, creating visible colonies in 2 days of incubation in the conditions described (*Methods*). In liquid medium, UTEX 3221 formed macroscopic aggregates of cells across all media and conditions tested, while UTEX 3222 exhibited unicellular, planktonic growth (Fig 1b, S1a). UTEX 3221 additionally exhibited phototactic motility, which appeared to be absent in UTEX 3222 (Fig S1b).

**Figure 1:**
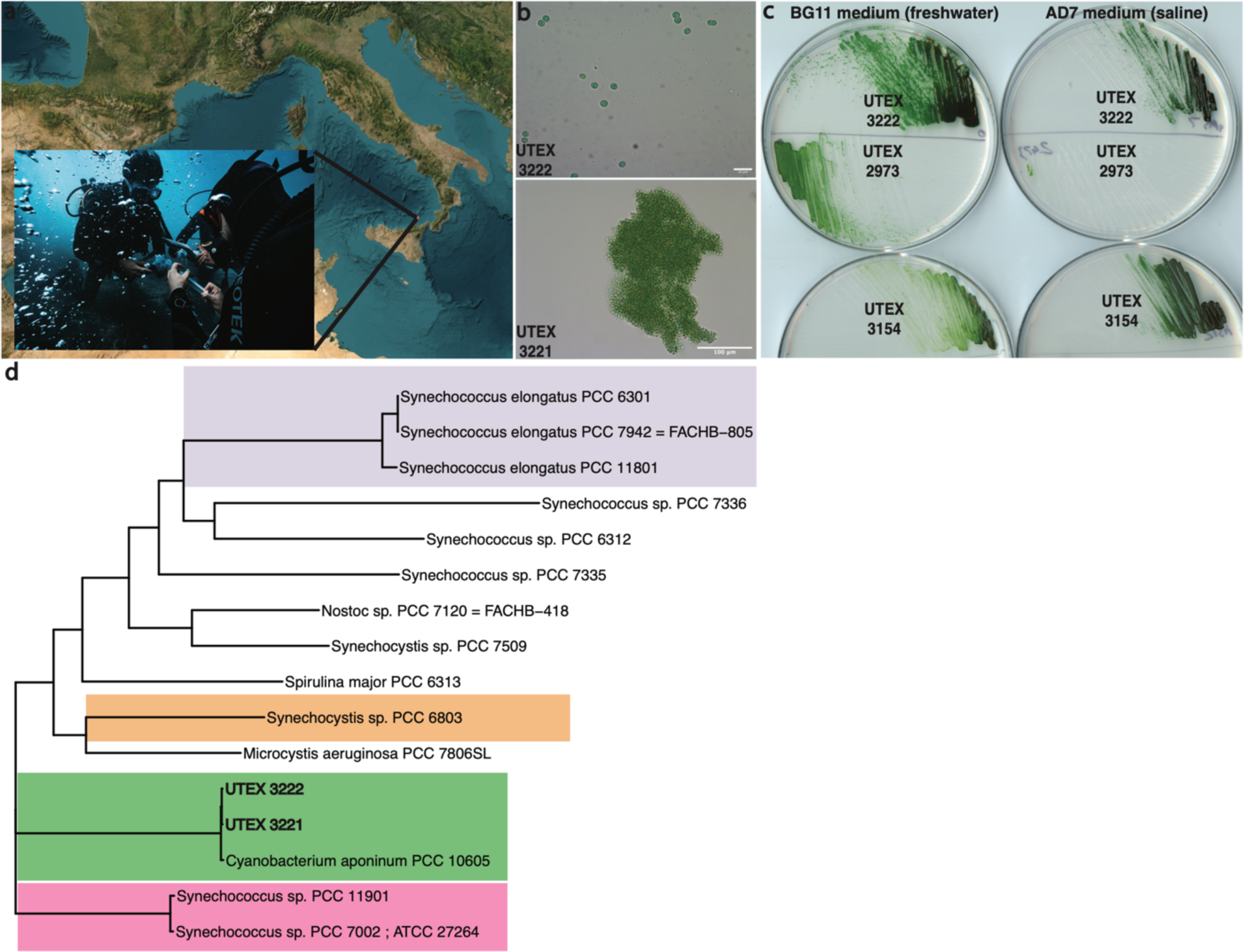
Isolation and sequencing. A) Samples were obtained from Baia di Levante, Vulcano, Italy. B) Micrographs of UTEX 3221 and UTEX 3222, displaying planktonic vs. aggregate growth. C) Axenic isolate UTEX 3222 growing alongside UTEX 2973 and UTEX 3154 (a derivative of PCC 11901) after 3 days at 37°C, 200µE, and 0.5% CO_2_. BG11 medium supplemented with vitamin B-12 D) Phylogenetic tree (using the Bac120 marker genes from the Genome Taxonomy Database) of novel isolates alongside their closest sequenced relative, and notable model cyanobacteria. Notable clades are highlighted in color

We chose to focus on UTEX 3222 for further characterization as planktonic growth is better explored in the existing literature. UTEX 3222 produced larger colonies than notable fast-growing cyanobacterial model strains UTEX 2973 or UTEX 3154 (PCC 11901 adapted to the absence of vitamin B-12)^19^ after 3 days of growth in the conditions tested (Fig. 1c, quantification in Fig S2), which did include 0.5% CO_2_. Notably, growth on either BG-11 freshwater or AD7 saline medium suggests that these strains are euryhaline. These isolates did not require vitamin B-12, which is required by PCC 7002, PCC 11901^18^, and to a lesser extent by UTEX 3154^19^. Both novel isolates exhibited an approximate 3.72±0.06 μm diameter spherical cell size, larger than described for other model cyanobacteria^12,18,29^.

### Genome Characteristics

Genome sequencing and assembly (see *Methods*) revealed approximately 4.6 MB genomes, and annotation identified coding regions, CRISPR elements, and other elements of interest (Table 1). UTEX 3221 and UTEX 3222 are closely related strains, sharing more than 98% Average Nucleotide Identity (ANI), and differing by at least two major genome inversion/translocation events (Fig S3). Phylogenetic comparison reveals the closest known sequenced relative of these strains to be *Cyanobacterium aponinum* PCC 10605 and confirms that these strains reside apart from clades containing highly-studied cyanobacterial model strains (Fig. 1C-E). Comparison of genomic regions by BLAST reveals genomic blocks with lower similarity among comparison strains, potentially highlighting novel elements in these genomes. antiSMASH^34^ identified biosynthetic clusters of interest, including terpenes, arylpolyene, lanthipeptides, and an iucA/iucC-like siderophore. Notably, PCC 10605 also exhibits these pathways.

**Table 1:** Summary of Genome characteristics. Annotations produced by annotation with BAKTA^30^, Prophage detection with PhiSPY^31^ using the PHROG database^32^, and Average Nucleotide Identity (ANI) using FastANI^33^.

|  | UTEX 3221 | UTEX 3222 | PCC 10605 |
| --- | --- | --- | --- |
| **Chromosome** | 4,417,179 bp | 4,280,321 bp | 4,114,099 bp |
| **Assembly Status** | circular | circular | circular |
| **Episomes** | None detected | None detected | 62,874bp circular |
| **tRNA** | 44 | 45 | 44 |
| **tmRNA** | 1 | 1 | 1 |
| **rRNA** | 9 | 9 | 9 |
| **ncRNA** | 9 | 9 | 9 |
| **ncRNA regions** | 5 | 5 | 5 |
| **CRISPR** | 7 | 6 | 11 |
| **CDS** | 3760 | 3602 | 3480 |
| **sORF** | 1 | 1 | 0 |
| **oriC** | 1 | 1 | 0 |
| **PhiSPY prophage candidates** | 2 | 2 | 1 |
| **ANI to PCC 10605 genome** | 98.41% | 98.45% | 100% |

Curiously, this close relative PCC 10605 was also isolated in Italy from thermal springs^35^, where studies have been conducted in regard to DNA replication^36^ and C-phycocyanin production^37^. Other relatives of PCC 10605 have been isolated worldwide, including from a marine environment near hot springs in China^38^, and a wastewater treatment system in Oman^39^. PCC 10605 did not appear to grow as quickly as these new isolates on solid medium in the conditions tested (Fig S4) and displayed an aggregation phenotype in liquid similar to UTEX 3221.

UTEX 3221 and UTEX 3222 thus display intriguing fast growth and genomic features and belong to a clade of cyanobacteria containing *Cyanobacterium aponinum* PCC10605 and other isolates. There is not extensive research on this clade, particularly concerning the investigation of fast-growing isolates and their application as chassis strains in synthetic biology. These strains have been deposited at the The University of Texas, Austin’s UTEX Culture Collection of Algae, and their genomes are accessible under UTEX 3221 and UTEX 3222.

### Growth characterization of UTEX 3222

To further characterize growth conditions for UTEX 3222, we measured the exponential growth rate across varying conditions in liquid culture. We began with growth in BG-11 medium, where UTEX 3222 had produced the most robust growth on petri plates (Fig 1B). In liquid medium, UTEX 3222 displayed a wide tolerance to temperature, with optimal growth rate at 45°C among the conditions tested (Fig 2A), somewhat higher than 30 or 37°C conditions used for most model cyanobacteria^14,18^. This temperature optimum is higher than previous reports in this clade of cyanobacteria^35,38^, and reflects realistic temperatures observed in outdoor photobioreactors midday in the absence of cooling equipment^40,41^. At the same time, the growth rate appears faster than thermophilic models like *T. elongatus* BP-1, which has an even higher temperature optimum of ∼57°C^42^. While noting this optimum at high temperatures, we continued at 37°C to facilitate comparison with pre-existing literature and isolates. No growth was observed at pH 5.5, but pH from 6.5 to 9.8 was well-tolerated (Fig 2B). The fastest exponential growth was observed at pH 6.5, identical to the isolation site, but higher growth density was quickly achieved at pH 8, likely due to better availability of bicarbonate at this pH (Fig S5B). Commensurate with its marine habitat, UTEX 3222 tolerated high levels of salt, even exceeding that of seawater, though a moderate 10 g/L NaCl produced the fastest growth (Fig 2C). Notably, this differs from results on solid medium, where BG-11 medium appeared to promote faster growth (Fig 1B). We did not observe elongated cells in elevated salinity, a phenotype occurring in the close relative PCC 10605^35^.

**Figure 2:**
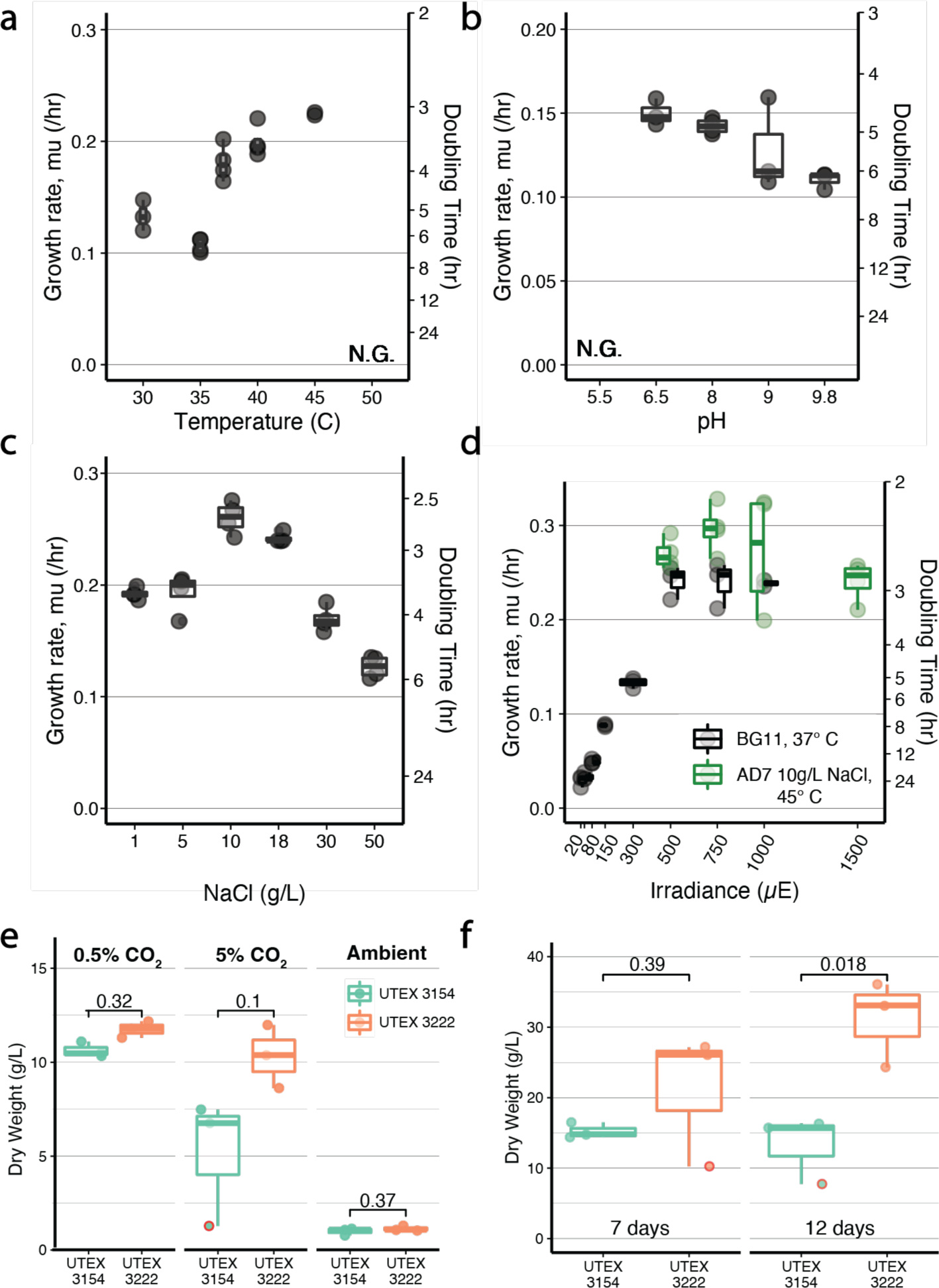
Growth conditions, and high-density growth. Exponential growth rate of UTEX 3222 in BG11 medium at varying Temperature (A), pH (B), Salt concentration (C), and total light (D). No growth (NG) was observed at 50°C, or at pH 5.5. Where otherwise not noted, growth conditions were BG11 medium at 37°C 500µE of light, pH 8.2, and 0.5% CO_2_. Experiments in panel C used AD7 medium modified to include the amount of salt shown (see methods for more detail). Modified AD7 was used in panel D for results in green, at 45°C. E) Biomass dry weights (gDW/L) after 7 day batch incubation in flasks, 200 µE light, 37C. F) Biomass Dry weight, after 7 and 12 days with light increased to 750µE on day 2, 1% CO_2_, 37°C. Triplicate experiments depicted with points, and additionally summarized by boxplots. Unpaired t-tests are depicted with brackets, with results either nonsignificant (ns) or significant (*, p<0.05). P-values reading left to right are 0.032, 0.1, 0.37 for B, and 0.39, 0.18, for C. Removing outlier low density cultures (circled in red) would result in P of .032, 0.068, .37, for E, and 0.00095, 0.049 for F.

UTEX 3222 tolerated irradiance of at least 1500 µE (Fig 2D), which is lethal for most photosynthetic microbes at these culture densities but tolerated by some fast-growing isolates^43^. In BG-11 medium at 37°C, growth rates increased with increasing light up to 500 µE, beyond which more light did not produce a higher growth rate. However, with results from the temperature and salinity tests in mind, we also tested AD7 medium with moderate salinity at 45°C. This condition supported an increasing growth rate up to 750 µE and promoted faster growth overall, with doubling times of 2.35±0.10 hours, the most favorable conditions tested (Fig 2D). It is likely that additional optimization could result in faster exponential growth, but we chose to instead focus next on high-density growth.

### High density growth of UTEX 3222

While fast growth on solid medium (Fig 1B) and reasonable growth rate at low density (Fig 2A-D) are good predictors of an organism’s facility in the lab and preferences among growth conditions, light-limited growth at high density is predicted to be more relevant for industrial applications, where high culture density is needed for high volumetric or areal productivity^44,45^. Following an initial observation of planktonic liquid growth of UTEX 3222 to high-density, we explored high density batch growth as a relevant industrial behavior.

Recent work reports record-setting cyanobacterial culture densities in PCC 11901^18^, and indeed, high-density growth appears to be a unique characteristic of this strain. Further development of PCC 11901 yielded UTEX 3154, a derivative with an alleviated vitamin B-12 requirement^19^. We thus compared directly to UTEX 3154, and used MAD2 saline medium previously optimized for high density growth^18^, to determine if UTEX 3222 shared the ability to grow to high density. Beginning with 0.5% CO_2_ and 200 µE light, we found that UTEX 3222 growth surpassed that of UTEX 3154, both by optical density over time (Fig S6A) and confirmed by biomass dry weight after 7 days (Fig 2E). Increasing CO_2_ to 5% surprisingly had a mildly negative impact on the dry weight of both strains, and the use of ambient air conditions (∼0.04% CO_2_) predictably led to far lower dry weight in both strains (Fig 2E). Increasing salt concentration appeared to improve dry weight, but not by a substantial margin for either strain and increasing the initial inoculum also did not meaningfully raise the biomass titer (Fig S6B-C). Increasing light to 750 µE after 1 day and increasing CO_2_ to 1% mirrors previous work^18^, and raised titers substantially (Fig 2F). In this condition, UTEX 3222 yielded 31.13±3.53 g/L biomass after 12 days, whereas UTEX 3154 yielded 13.25±2.76 g/L. Collectively, the findings suggest that UTEX 3222 demonstrates a growth phenotype of high density, rivaling and even surpassing what has been reported for the PCC 11901 and UTEX 3154 strains^18^. Further work to optimize medium and conditions specifically for UTEX 3222 could increase yield further.

### Biomass composition of UTEX 3222

Because UTEX 3222 is not closely related to well-studied model cyanobacteria (Fig 1C), we expected its cellular composition to differ. These differences could inform efforts to use the cyanobacterial biomass itself or potentially engineer this organism to produce new products. We began by exploring cellular composition qualitatively by Transmission Electron Microscopy (TEM) of cells from high-density culture, revealing putative extracellular polysaccharides (EPS), as well as what appeared to be storage granules for glycogen or polyhydroxyalkanoates, both common carbon storage products in cyanobacteria(Fig 3A, S7A,B,D)^46^. Relevant biosynthetic genes for both products are found in the UTEX 3222 genome. Further images reveal heterogeneity; cells grown in the conditions tested appear to vary in the number and size of storage granules (Fig S7). UTEX 3154 grown and imaged in a similar manner did not display as prominent EPS or storage granules (Fig S7E,F).

**Figure 3:**
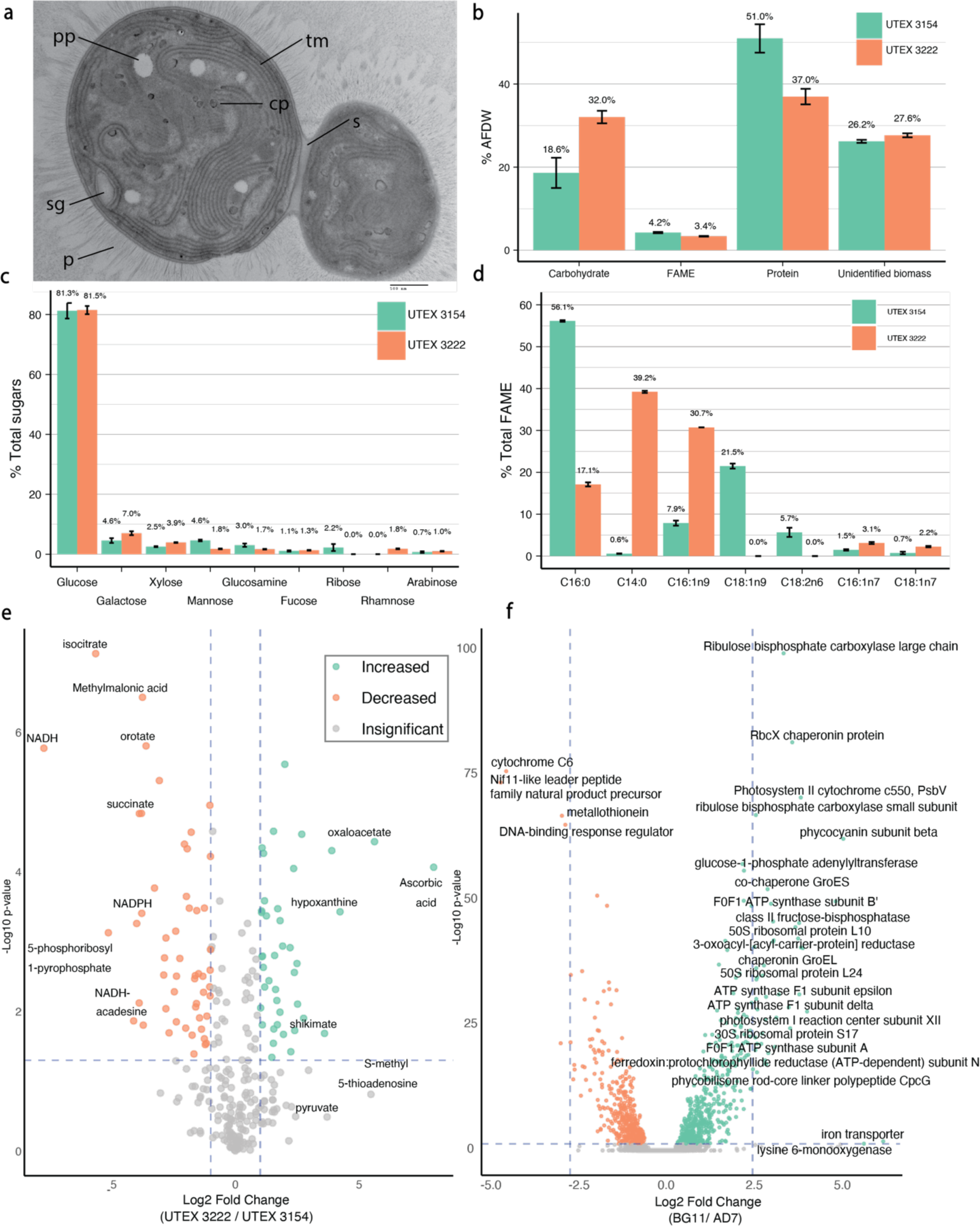
Characterizing UTEX 3222 biomass. A) Representative TEM image, with abundant Thylakoid membrane (tm), division septa (s), and putative polyphosphate granules (pp), storage granules (sg), cyanophycin granules (cg), and extracellular pili (p). B) Major macromolecule composition, as a percentage of Ash-Free Dry Weight (%AFDW). C) Sugars detected following acid hydrolysis of biomass. D) Fatty Acid Methyl Ester (FAME) species, species with a mean measurement >2% shown, for lower abundance species see Supplemental Figure 9. For all charts in this figure, bars depict the mean of triplicate culture measurements, and error bars depict the standard error of these measurements. E) Relative comparison of a panel of polar metabolites across UTEX 3222 and UTEX 3154 biomass grown to high density. Analytes differing in abundance by more than 10-fold are labeled with text. F) Differential expression analysis by RNA-sequencing of UTEX 3222, comparing freshwater (BG-11) growth to saline (AD7) medium.

Quantitative analysis of overall biomass composition revealed a higher proportion of carbohydrates in high-density UTEX 3222 biomass compared to UTEX 3154, with correspondingly lower fractions of ash-free dry weight consisting of protein and other components (Fig 4B, Fig S8A). This overall difference was also observed in C/H/N elemental analysis, with UTEX 3222 biomass having higher overall carbon content (Fig S8B). These measurements argue that UTEX 3222 may produce more storage carbohydrates than UTEX 3154 when grown to high density. This could explain in part differences in the biomass density they achieve in the same conditions.

**Figure 4:**
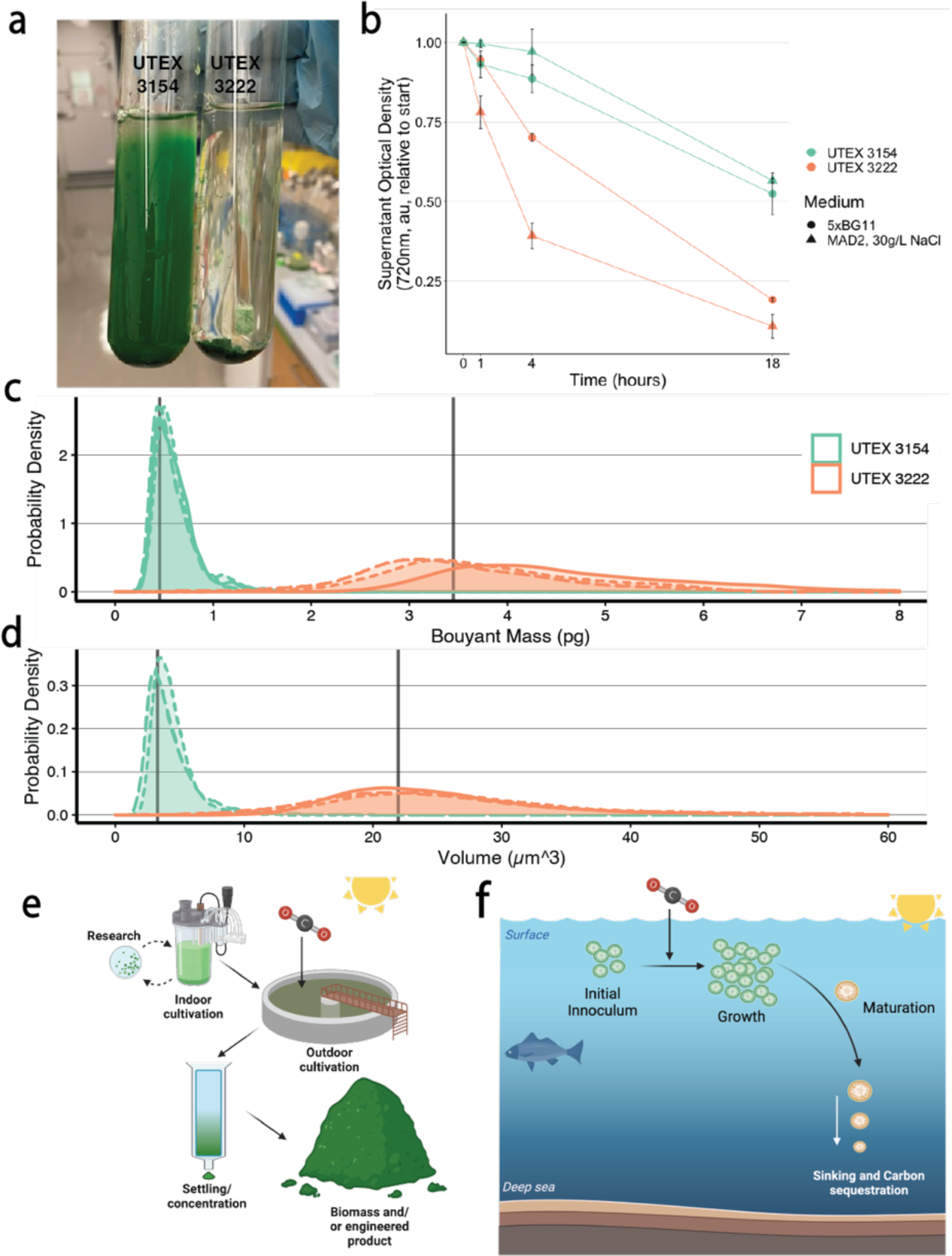
Sinking behavior of UTEX 3222. A) Photograph of cultures allowed to settle for 12 hours at 20°C, after 12 hours of growth at 37°C in BG11 medium. B) Sinking timecourse, employed with high-density batch-cultivated cultures at 20°C in the absence of light. Lines and points display the mean of triplicate experiments, error bars are standard error. C) Buoyant masses of single cells, as measured using the SMR (see methods). D) Volumes of single cells, as measured using the CC (see methods). Data in C and D is presented as a probability density function with individual replicate experiments plotted separately with different line styles. In C, the difference between replicates reflects biological variance in cells’ buoyant density. The peak of this density function approximates the mode of data, and the mean of these peaks is shown. E) Schematic summarizing that these strains support rapid research and development due to their fast growth rates in the lab, and that their facile sinking phenotypes could be leveraged to improve production of biomass and/or engineered products at scale. F) Schematic summarizing a potential marine carbon sequestration procedure, in which sinking phytoplankton preferentially support movement of carbon into deep ocean sediments.

Acid hydrolysis of carbohydrates derived from either strain reveals the overall composition of component sugars, with >80% of carbohydrates in both strains digesting to glucose but significant differences in lower-abundance sugars (Fig 4C). In particular, where rhamnose comprises nearly 2% of hydrolyzed sugars from UTEX 3222, it is undetectable in UTEX 3154, and ribose was not detected in UTEX 3222. Notably, both strains contain relatively low concentrations of lipids as quantified by fatty acid methyl esters (FAME, Fig 4B, Fig S9), which can be quite abundant in oleaginous model phototrophs. The differing composition of these FAME across the two strains was notable; however, UTEX 3154 biomass contained far more palmitic acid (C16:0) and abundant oleic acid (C18:1n9) and linoleic acid (C18:2n6), which were both undetectable in UTEX 3222 (Fig. 4D). Conversely, the FAME in UTEX 3222 contained far more myristic acid (C14:0) and hypogeic acid (C16:1n9) (Fig. 4D), and had overall shorter FAME chain lengths.

Similarly, metabolites can be compared between these strains in order to help understand their differences and characterize UTEX 3222 as a chassis for metabolic engineering and as a source of high-density phototrophic biomass. Mass spectroscopy of high-density biomass of UTEX 3222 and UTEX 3154 provides relative quantification of metabolites, and by comparing a targeted panel of 292 common polar metabolites, we see myriad differences emerge (Fig 3E). UTEX 3222 produces ascorbic acid (vitamin C), which is undetectable in UTEX 3154. Shikimate, a precursor in the biosynthesis of aromatic compounds, is elevated compared to UTEX 3154. Similarly, mevalonate, a precursor for terpene synthesis, is elevated in comparison to UTEX 3154. These differences were observed when growing strains in identical medium and conditions; further characterization across multiple conditions and physiological states would provide additional context. For instance, NADPH and NADH are more abundant in UTEX 3154, but these energy carriers are more likely to change quickly depending on the physiological state.

Transcriptomics can help describe physiological differences across media conditions, assist with genome annotation, and identify regulatory elements. We thus interrogated the transcriptome of UTEX3222 when grown in either AD7 saline medium or BG11 freshwater medium by strand-specific RNA sequencing. As expected, the varying media types yielded dramatically different expression profiles. 1,026 genes were significantly (adjusted p-value < 0.05) increased in expression in BG11, whereas 948 increased in AD7 medium (Fig 3F). This corresponds to 52% of coding frames in the genome, significant variation in expression. Overall, genes associated with growth and central metabolism (e.g., RuBisCO, light harvesting proteins) were increased in expression in BG11. Unexpectedly, we identified potential responses to iron starvation in BG11, and in nitrogen starvation in AD7 medium, highlighting that these common media might limit growth, even for low-density batch cultures. Overall, transcriptomics proved valuable for understanding response to varying medium conditions and suggesting modifications to growth medium. The strand-specific alignments produced can additionally refine our genome annotation, yielding clues about expression level, promoter elements, operons, etc.

### Sinking/settling phenotype

In the routine handling of UTEX 3222, we observed that liquid cultures were prone to settle after several hours without agitation, with essentially all biomass concentrated in a tight pellet, whereas cultures of other fast-growing strains did not settle as quickly nor completely (Fig 4A). This differing behavior could potentially be of use for industrial processing, where concentration and dewatering of biomass is a substantial economic challenge^47,48^, estimated to contribute from 15% of production cost^49^, to as much as 30% of production costs^50^. A time course provided a more detailed look at cell settling, confirming that high-density UTEX 3222 biomass settles more quickly in comparison with UTEX 3154 in an artificial seawater medium (Fig 4B). This is the case whether the high-density culture was grown in saline MAD2 medium or in freshwater 5X BG11 medium, although curiously, culture grown in saline medium appears to settle more quickly (Fig 4B).

Given UTEX 3222’s lack of phototactic motility, two differing phenomena could drive this settling behavior-aggregation of single cells into larger particles that sink more readily and/or individual cells having faster settling rates. Aggregation plays a key role in the behavior of fast-settling mutants of PCC 7942 cyanobacteria^51^, but can be predicated on the ionic strength of the settling medium and can be mimicked with chemical flocculants^49^. We were curious whether individual cells had advantageous settling behavior, because this could drive the sinking of biomass in more dilute conditions, such as cells growing in marine habitats and affecting natural carbon cycles^52^. The gravitational sinking of a cell, as defined by Stokes’ law, is dependent on cell volume and buoyant density. We measured the buoyant masses and volumes of single cells as previously described (Fig 4C-D)^52^. As buoyant mass is a function of cell volume and cell buoyant density, this allows us to solve for the buoyant densities and, thereby, gravitational sinking velocities of the cells. This revealed that individual cells of UTEX 3222 have 2.16-fold faster gravitational sinking velocities than UTEX 3154 comparison cells (Table 2). This difference in sinking velocities was mediated by cell volume differences rather than buoyant density differences. Notably, one replicate culture of UTEX 3222 displayed an increased buoyant mass without a difference in volume, indicating higher buoyant density and ∼28% faster gravitational sinking velocity. Previous results point to starvation responses potentially increasing algal mass and sinking velocity^52^, which is in line with the putative storage polymers observed in UTEX 3222 (Fig S8) and the putative nitrogen starvation response observed in this growth medium (Fig 3F).

**Table 2:**
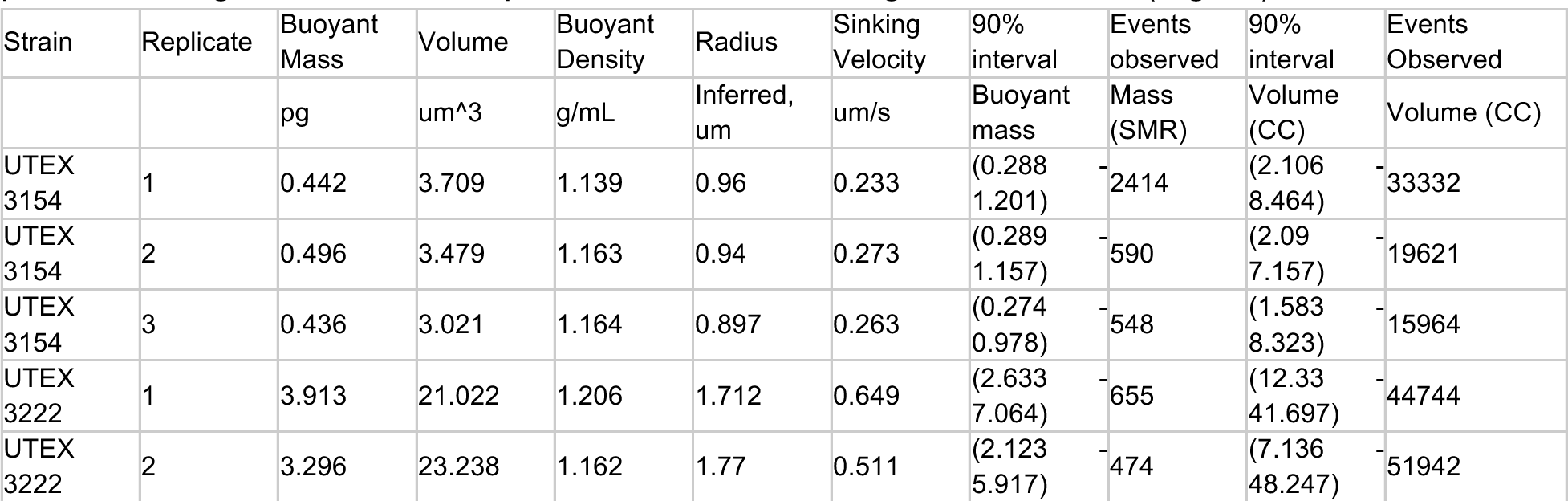
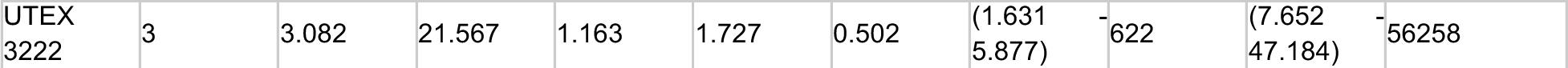
Physical parameters of UTEX 3222 cells drive different sinking behavior. Buoyant masses of cells were measured with a Suspended Microchannel Resonator (SMR), and volumes were measured with a Microsizer, or “Coulter Counter” (CC) instrument (See Methods, Fig 4C-D). Summary of data are reported alongside the intervals in which 90% of the data is contained, and the number of observations made.

While UTEX 3222’s gravitational sinking velocity differs by a significant margin from UTEX 3154, the predicted rates are insufficient to account for the stark difference in settling behavior fully (Fig 4A,B). This indicates that aggregation or other factors are likely significant drivers of this difference, as well. Notably, the sister strain UTEX 3221 and the relative strain PCC 10605 exhibit large aggregates in the conditions tested that settle far more quickly (Fig S1), and may offer clues for further manipulating aggregation and settling, offering additional industrial benefit. A better understanding of phytoplankton sinking in seawater also improves our understanding of these organisms’ involvement in natural carbon cycles^52,53^, and presents intriguing possibilities wherein sinking behavior could be manipulated to drive stable carbon sequestration in the deep ocean (Fig 4F).

## Discussion

In this study, we identified, sequenced, and characterized UTEX 3222 and UTEX 3221: two photosynthetic fast-growing strains of *Cyanobacterium aponinum* isolated from high CO_2_ marine volcanic seeps in the coastal Mediterranean Sea. UTEX 3221 formed cell aggregates during growth, whereas UTEX 3222 displayed rapid, planktonic growth to high density; given this result, we chose to further investigate UTEX 3222’s potential as a biotechnological chassis. We propose that 1) its rapid growth on solid medium in the lab, out-performing comparison strains UTEX 2973 and UTEX 3154 makes it an intriguing, facile lab model 2) Its rapid growth in liquid culture (as fast as 2.35 hr doubling time in this study), and tolerance of high light, high salinity, and high pH could support biotechnological applications 3) Its growth to exceptionally high density in batch culture (>30 g/L dry weight, >2x that of the relevant comparison strain in this work, and higher than existing reports to our knowledge) could benefit for productivity and harvesting, 4) the composition of its biomass differs from existing model strains, possibly presenting new opportunities for biomass valorization and metabolic engineering, and 5) the sinking/settling behavior of this biomass offers potential benefits for biomass harvesting, or toward sequestration of carbon in marine environments.

It is virtually impossible that the ideal host for photosynthetic bioproduction has already been isolated. This fact alone should motivate further efforts to isolate and characterize new strains from a variety of environments. When comparing strains, it can be difficult to make an apt comparison. For instance, UTEX 2973 is understood to be the fastest-doubling cyanobacterium isolated to date. In this work we not only observe UTEX 3222 producing superior growth on solid medium (Fig. 1B), but also a growth rate 45% faster than that reported for UTEX 2973 when grown in similar light and CO_2_ conditions in the same instrument (Fig 2D) (https://doi.org/10.1128/spectrum.00500-23). This result is promising but should be viewed in a broader context, wherein UTEX 2973 is also capable of exemplary exponential growth when growth is optimized with very high light and CO_2_ conditions (1.5-hour doubling, ∼57% faster growth rate)^14^. Faster growth rates for other strains have been reported, albeit in differing conditions and methods of measurement^18^. Growth rate measurement in cyanobacteria is also difficult to reproduce between labs, varying as much as 36% even when the same strain and methods are used, likely due to the intrinsic sensitivity to light quality and air composition^54^. Ultimately we focused on high-density growth, where UTEX 3222 out-performed the comparison strain UTEX 3154 by more than 2-fold greater biomass titer after 12 days, which is a potentially more biotechnologically relevant trait than exponential growth rate^44,45^.

Marine phytoplankton accounts for about half the photosynthetic primary production on earth (165-183Gt CO_2_/yr)^72,73^, fixing approximately 3-fold as much carbon as total anthropogenic greenhouse gas emissions(59±6.6 CO_2_e/yr)^74^. It’s estimated that about one-fifth of this carbon captured is exported to the deep ocean^75^. Thus, approaches that could meaningfully increase this fraction could have a tremendous impact and are an area of active study^76^. The Intergovernmental Panel on Climate Change (IPCC), as well as the National Academies of Sciences, have highlighted the need for negative emissions technologies (or "carbon sequestration") to avoid the worst effects of anthropogenic climate change^69,77^. These same bodies also acknowledge that we suffer from a shortage of proven solutions to fit this need and recommend developing a variety of approaches to achieve negative emissions. The cyanobacteria isolated here show potential to help solve longstanding challenges in this area. By accumulating carbon-rich storage polymers internally and carbon-rich EPS externally, such strains could accumulate a very high carbon-to-nutrient ratio. This offers a potential solution to the “nutrient-robbing” problem plaguing the marine biological pump, wherein precious nutrients are co-sequestered along with the carbon contained in phytoplankton biomass^76^. In addition, because the most abundant organisms in the ocean are very small cyanobacteria^78^, which are expected to have low sinking rates, interventions that shift marine microbial populations toward larger, faster-sinking organisms would be expected to increase the fraction of carbon exported to the deep ocean, rather than cycling back into the atmosphere. The strains described here accumulate carbon-rich storage polymers and sink readily in seawater. Their fast growth in lab conditions facilitates further experimentation toward the development of these kinds of interventions.

In total, our core hypothesis was that shallow carbon dioxide seeps would contain novel organisms for carbon sequestration, and our results indicate that is the case. Therefore, while UTEX 3221 and UTEX 3222 are promising biotechnological chassis, we claim that their isolation only further indicates the potential of searching natural and underexplored environments for novel microbial life and diversity (which in turn mandates conservation of these ecosystem). Further, culturing rather than sequencing provides advantages in detecting rare organisms below the detection limit of metagenomic sequencing and providing a detailed understanding of the behaviors of specific organisms. Overall, we expect that continued microbial exploration of CO_2_ seeps and other interesting ecosystems will yield further, more optimal organisms for carbon sequestration and other societally important challenges.

## Methods

### Sampling and Isolation

Hundreds of samples of seawater and sediment were collected along a well-established pH/*p*CO_2_ gradient in Baia di Levante (Vulcano Island, Italy). To assess the spatial variation in the carbonate chemistry, a Hobo® MX2501 submersible pH/Temperature logger and a Hydro II CO_2_ logger (Contros System & Solutions GmbH, Germany) were deployed at 1-4 m depth in each sampling site. Seawater pH (NBS scale) and temperature (T, °C) were recorded at 1 minute intervals. The pH logger was calibrated with standard buffer solutions (for NBS scale) of pH 4.01, 7.00 and 10.00, and then converted to Total scale (pHT). The Hydro II CO_2_ logger recorded pCO_2_ (μatm) every 10 sec. Seawater samples for Salinity and Total alkalinity (TA) were also collected in triplicate. Seawater was filtered at 0.45 μm using disposable cellulose acetate filters and stored at room temperature in the dark until TA was measured by a titration system (TiTouch i915, Metrohm). The titrations were cross-validated using a working standard (SD: ± 9 μmol kg-1) and against certified reference material from the A.G. Dickson laboratory. 50mL samples were obtained using syringes and kept at ambient temperature in 50mL conical tubes with exposure to light for several days, then packed and shipped in darkness at approximately 4°C. Seawater pCO_2_ was calculated from pHT, temperature (T; °C), salinity and total alkalinity (TA; mmol kg-1) using the Carb function (flag = 8) in the seacarb package (Lavigne and Gattuso, 2010) in RStudio software (version 4.2.1)<u>(Seawater Carbonate Chemistry [R packa…; Team and Others 2011)</u>.

Multiple samples per dive site were pooled by filtering using a sterile 0.22μM filter and then rinsing material from the filter using 50mL of Enrichment medium. Enrichment medium (AD7++) was AD7 medium prepared as per Włodarczyk et al.^18^, and modified by including 100mg/L of cycloheximide to inhibit the growth of eukaryotes, addition of ATCC trace mineral supplement (MD-TMS, ATCC) and ATCC vitamin supplement (MD-VS, ATCC) at 1/200th strength, and substitution of Tris-HCl for 10mM of TES-KOH pH 8.2. After initial enrichment, routine culture was performed in AD7 medium or BG-11 medium, also buffered with TES-KOH pH 8.2. Enrichment cultures were cultivated in 250mL baffled glass Erlenmeyer flasks, closed by both foam stoppers and clear plastic flask closures. Cultures were incubated at 0.5% CO_2_, with 200μE of white LED light at 37°C in an illuminated incubator (Multitron, InforsHT) shaking at 220rpm. For some enrichments, visible white growth occurred after 1-2 days. This growth was likely from Iron-oxidizing bacteria due to Iron present in sediment samples. Such enrichments were subjected to a further 100-fold dilution in AD7++, and in some cases resulted in green, photosynthetic unicellular growth after 6 days in these conditions. Some enrichments yielded macroscopic green "spidery" growth, which appeared to be filamentous cyanobacteria, which proved difficult to render axenic.

Samples from two enrichments exhibiting unicellular green growth were streaked on plates of AD7++ solidified with 1% agar (bacteriological grade, APEX). These were incubated at 0.5% CO_2_, with 200μE of white LED light at 37°C in a growth chamber (AL-41L4, Percival). Green colonies were observed alongside white colonies, presumed to be heterotrophs subsisting on agar, buffer, or cyanobacterial exudate. Green colonies were restreaked to solid AD7++ after 3-4 days, and this procedure was repeated at least 5 times with the goal of obtaining axenic cyanobacterial cultures. Multiple isolates from each enrichment were rendered axenic, but in all cases, these were later determined to be identical strains. Filamentous cyanobacteria were plated to solid medium, but proved difficult to separate from non-photosynthetic contaminants, and were excluded from further characterization. Photos on solid medium were obtained on a flat-bed scanner (Epson) and colony size was measured by image analysis in Fiji^79^.

### Culture and cryo-preservation

Routine growth of liquid cultures was performed by culturing a 10mL volume in 50mL Erlenmeyer flasks in AD7 or BG-11 medium, in conditions as above. Cryo-preservation was performed by the addition of DMSO to 9%, flash-freezing with liquid nitrogen, and storing at -80°C. Frozen cultures are revived by streaking to AD7 or BG-11 medium and incubating as above, but with <100μE light and ambient air for the first day, and then as above for subsequent days. Cryo-preserved cultures remain viable in this way for more than one year and likely much longer. Incubator temperatures were confirmed with a NIST-traceable thermometer (Digi-Sense), and photosynthetically active light was confirmed with a light meter (MQ-500, Apogee).

Light microscopy of wet-mounted liquid cultures was performed using a Zeiss Axio Imager Z-1 microscope, illuminated by a white LED light source. Phototactic motility was investigated by adapting published methods^80^. Briefly, liquid culture was plated on BG11 with 0.3% agar (“swim medium”), and plates were incubated in a foil packet to ensure light only enters from one side. These cultures were incubated for 4 days in conditions as above. Limited light available inside the foil packet likely resulted in slower growth overall.

### Genome Sequencing and Analysis

Cultures were grown as above, and 5mL of overnight culture was pelleted at 4 krcf and stored at -20°C. Light microscopy (Fig. 1B, S1) and lack of observed growth on LB medium suggested isolates were likely axenic, an assertion supported by the lack of contaminating sub-assemblies after DNA assembly. DNA was extracted from frozen pellets using the ZymoBIOMICS DNA Miniprep Kit (Zymo, Cat. # R2002), performing the initial bead disruption step using a Tissuelyzer LT (Qiagen, Cat. # 85600) set to 50Hz for 30 minutes, at 4°C. DNA was quantified using the Qubit 1xds DNA, broad range assay (ThermoFisher Scientific, Cat. # Q33265). Long read sequencing was performed by Plasmidsaurus (Eugene, OR) or SeqCenter (Pittsburgh, PA), using V14 ligation-based library prep (Oxford Nanopore Technologies, Cat. # SQK-LSK114) and R10.4.1 flow cells (Oxford Nanopore Technologies, Cat. # FLO-PRO114M) on the PromethION device. Short-read genome sequencing was performed by SeqCenter using the Illumina tagmentation DNA prep kit (Illumina) and sequenced on a Novaseq X instrument (Illumina) performing a 2X151bp run, using custom 10bp indices. Demultiplexing, adapter trimming, and quality control was performed using bcl-convert v4.1.5 (Illumina).

Reference genomes were constructed using long nanopore reads by assembly using Flye^81^, and consensus polishing using Medaka (Oxford Nanopore). These genomes were subjected to further short read polishing using PolyPolish^82^ and annotation using Bakta^30^ via the Bakta web tool. Additional assembly was performed by two methods: hybrid assembly in Unicycler^83^ followed by Circlator^84^, and hybrid assembly in Trycycler^85^, to attempt to detect episomes that may have been excluded from the Flye assembly, but none were detected. Prophages were predicted using PhiSPY^31^, supplying the PHROG database^32^ for Hidden Markov Model comparison. Evolved mutants were sequenced using Illumina short read sequencing as described above, and resulting reads were compared to reference using BREseq^86^. Visualization of genomes was performed using ProkSEE^87^ via the Proksee web tool, including analysis using the CARD RGI, CRISPR/Cas Finder, Alien Hunter, mobileOG-db, Phigaro and Virsorter, and FastANI^33^ plugins. Annotated genomes were routinely viewed using Geneious Prime software (Dotmatics). Genomes were scanned for biosynthetic clusters using antiSMASH 7.0^34^.

### Culturing and Growth Rate measurement

To measure exponential growth rate in liquid, a Multicultivator instrument with light upgrade was used (MC-2500-OD, Photon Systems Instruments). In this system, air and CO_2_ were mixed using a Gas Mixing System (GMS-150, Photon Systems Instruments), and this mixture was first humidified by bubbling through water, then sparged into the cultures using a 5 X 210mm porosity B glass filter stick (Ace Glass).

Liquid cultures were prepared in shake flasks as described above, then inoculated into 50mL of relevant media in Multicultivator vials at an OD_720_ of 0.1, as measured in cuvette using a Nanodrop 2000 (Thermo Scientific). These cultures were first acclimated for 1 hour at 100µE light at growth temperature, then light was increased and OD measurement initiated. Unless otherwise stated, cultures were grown in BG-11 medium with 4µg/L Vitamin-B12 (to facilitate comparison with B-12 auxotrophs), at 37°C, 0.5% CO_2_ flowing at 0.1 L/min into each vial, and 500µE of light. Growth rate µ was inferred by fitting an exponential curve to optical density values measured by the Multicultivator between 0.1 and 0.35 and interpreting the slope of the fit using a custom R script (see figure S5A for representative fits). Doubling time was calculated as ln(2)/µ.

### High Density Batch Growth

High density batch growth cultures were grown in MAD2 medium prepared as per Włodarczyk et al.^18^, and modified by substitution of Tris-HCl for 10mM of TES-KOH pH 8.2. Some precipitation is observed. 2uL of Antifoam 204 is added to each flask, as a 20% solution in 70% ethanol. Patches of relevant strains on solid medium were inoculated into 10mL AD7 medium and incubated for ∼20h as above. The resulting culture was used to inoculate 50mL of MAD2 medium to an OD720 of 0.1 in a 250mL baffled shake flask fitted with sponge top and plastic closure. Cultures were incubated as indicated, with the 200µE condition in an Infors-HT incubator and the 750µE condition in a Percival incubator. For the 750µE condition, only 200µE of light was used for the first 24h of incubation, and 750µE was used thereafter. At the conclusion of growth, samples were pelleted in preweighed 50mL conical tubes. Dry weight was determined after 48 hour lyophilization (Labconco FreeZone).

### Microscopy

Light microscopy was performed by imaging a wet mount of liquid culture using a Zeiss Axio Imager Z1. Images were processed using FIJI software. Transmission Electron Microscopy (TEM) images were processed by Harvard Medical School EM facility. TEM samples were fixed in a solution of 1.25% formaldehyde, 2.5% glutaraldehyde, and 0.03% Picric acid in 0.1M cacodylate buffer, pH 7.4. Fixed samples were stained with Osmium tetroxide and uranyl acetate, then dehydrated using an ethanol series followed by propylene oxide. Samples were infiltrated with a 1:1 mixture of EPON resin (Westlake) with propylene oxide for 16 hours at 4°C, then polymerized in Epon resin for 24 hours at 60°C. Embedded samples were sectioned by standard methods before viewing using a JEOL 1200EX transmission electron microscope.

### Biomass characterization

Biomass composition was characterized at the National Renewable Energy Lab (Golden, CO) as reported previously^88^. In brief, an Elementar VarioEL cube CHN analyzer was used to determine the C/H/N content. A multiplication factor of 4.78 was used to estimate the total protein content from N content^89^. Samples were subjected to acid hydrolysis and the resulting monomeric sugars measured with a Carbopac HPAEC-PAD system with PA-1 column. Where a sugar was undetectable, a value of 0% was used. Where a sugar was below the limit of quantification, a value 1/5th the mean of other measurements for that sugar was substituted.

### Metabolomics

Sample preparation was adapted from Jugder et al^90^. Briefly, 400uL of culture was harvested by centrifugation, then subjected to extraction in 800uL 80% Methanol for 20 minutes in an ultrasonicator bath (Elmasonic P, Elma) at 20°C. Cell debris was removed by centrifugation, and the supernatant was incubated at -80°C for 16 hours and centrifuged again. The resulting supernatant was dried in a SpeedVac Vacuum Concentrator (Thermo) under vacuum and ambient temperature, then stored at -20°C. Samples were analyzed by the Beth Israel Deaconess Medical Center Mass Spectroscopy Core Facility as per previous studies^90^ using a 6500 Qtrap triple quadrupole mass spectrometer (AB/SCIEX), Prominence UFLC HPLC (Shimadzu), and selected reaction monitoring (SRM) of 298 water-soluble metabolites using MultiQuant v3.0 software (AB/SCIEX). The resulting peak areas (Supplemental Data 6) were analyzed and visualized using Metaboanalyst.ca, using normalization to median, log10 transformation, and mean-centering functions.

### RNA-sequencing and downstream analysis

10mL cultures were grown in 50mL unbaffled flasks, at 37°C and 200µE light, shaken at 220 RPM in a growth chamber maintaining 0.5% CO_2_ for 16 hours. Four replicate cultures were prepared in either BG11 medium or AD7 medium. RNA was isolated from using the Monarch Total RNA miniprep kit (NEB), using the Tough-to-lyse sample protocol with bead-beating (Tissuelyser LT, Qiagen). Ribosomal RNA depletion and strand-specific RNA-sequencing prep was carried out by Azenta Biosciences, along with 2X150bp paired end sequencing on a HiSeq instrument (Illumina). Demultiplexing using bcl-convert (Illumina) and 8bp index pairs resulted in >40 million read pairs for each sample. Sickle (https://github.com/najoshi/sickle) was used in paired-end mode to quality filtering (minimum quality = 35, minimum length = 45) and trim adapters from raw sequencing data. We aligned filtered reads with Bowtie2^91^ (params: --very-sensitive-local) to the open reading frames predicted in our annotated genome for UTEX 3222. DEseq2^92^ processed these raw alignment counts to compute differential abundance and statistical metrics. We report significant findings as those with an adjusted p-value of less than 0.05.

#### Measuring settling phenotypes and cell physical parameters

Sinking/settling of cells was quantified by filling a polystyrene cuvette (Brand-Tech, Semi-micro 759076) with 1mL of culture diluted to approximately OD_720_ of 1 with culture medium, and measuring OD_720_ over time using a Nanodrop 2000c instrument (Thermo). Because this instrument’s light path is 8.5mm above the bottom of the cuvette, it provides a measure of the OD_720_ above a pellet that accrues over time. Settling occurred at 20C, shielded from light.

Buoyant masses of single cells were measured using a Suspended Microchannel Resonator (SMR), where single cells flow through a microfluidic channel inside a vibrating cantilever, and the change in vibrational frequency of the cantilever is measured and converted to buoyant mass^93,94^. In this study, the SMR had a cantilever cross-section of 8×8 µm, and the resulting frequency measurements were interpreted by custom Matlab code, as previously demonstrated^95^. Volumes of single cells were measured using Multisizer 4 Instrument (Beckman Coulter, also known as Coulter Counter, or CC). The SMR and CC were pre-filled with AD7 medium and measurements were performed at approximately 20°C. Both measurements were calibrated using NIST traceable polystyrene beads (Thermo Scientific, Duke Standards). Cultures for these experiments were grown in AD7 medium with 0.5% CO_2_ and 200 µE light at 37°C, for approximately 16 hours. Triplicate measurements were made across unique cultures and multiple days. Gravitational cell sinking velocities were derived as previously reported^52^, using Stokes’ Law. For solving Stokes’ Law, we assumed a spherical cell shape, a medium density of 1.02 g/mL, and a dynamic viscosity of 1.07E-3 Pa*s. Cell radius was calculated from cell volume measurements and buoyant density was calculated from cell volume and buoyant mass measurements. To minimize the influence that population outliers (e.g., cell aggregates) have on the results, we used the modal buoyant mass and volume values, as determined using probability density functions.

## Funding and acknowledgements

We thank the Pakrasi lab for their input and guidance in cyanobacterial research. We additionally thank the Harvard Electron Microscopy facility and the Beth Israel Deaconess Mass Spectrometry Core Facility for their contributions to this work. Funding was provided by the US Department of Energy (DOE) under grant no. DE-FG02-02ER63445 and by the National Science Foundation (NSF) award no. MCB-2037995 (both to G.M.C.), as well as by a contribution from SEED Labs, a division of SEED Health, Inc. Seed Health has no intellectual property or other ownership over the strains described in this study. We additionally thank the WorldQuant Foundation for incidental funding and the Scientific Computing Unit (SCU) at Weill Cornell Medical College. We are also grateful to Prof. Alessandro Aiuppa (University of Palermo) for his useful contribution, Dr Francesco Italiano and Dr Alessandro Gattuso (INGV, Italy) for helping with field logistics, the Diving Centre Saracen at Marina di Vulcanello and the local community of Vulcano Island for their support. MM and DS were also supported by the International CO_2_ Natural Analogues (ICONA) Network.

## Data Availability

Strains are publicly available through the UTEX algae culture collection. Raw sequence data and assemblies are available at BioProject PRJNA1033390.

## Author Contributions

MGS, JRH, and BT conceived of the study with input from TC, PQ, MM, CEM, and GMC. KR, JRH, DS, GT, PQ, MM, and BT organized the Vulcano expedition and performed sampling and sample preparation. MGS performed isolation and characterizations of strains and analysis of data, with help and input from TCT, IGM, and TC. MGS prepared initial drafts of the manuscript. All authors contributed to editing the manuscript.

## Supplemental Figures

**Supplemental Figure 1.**
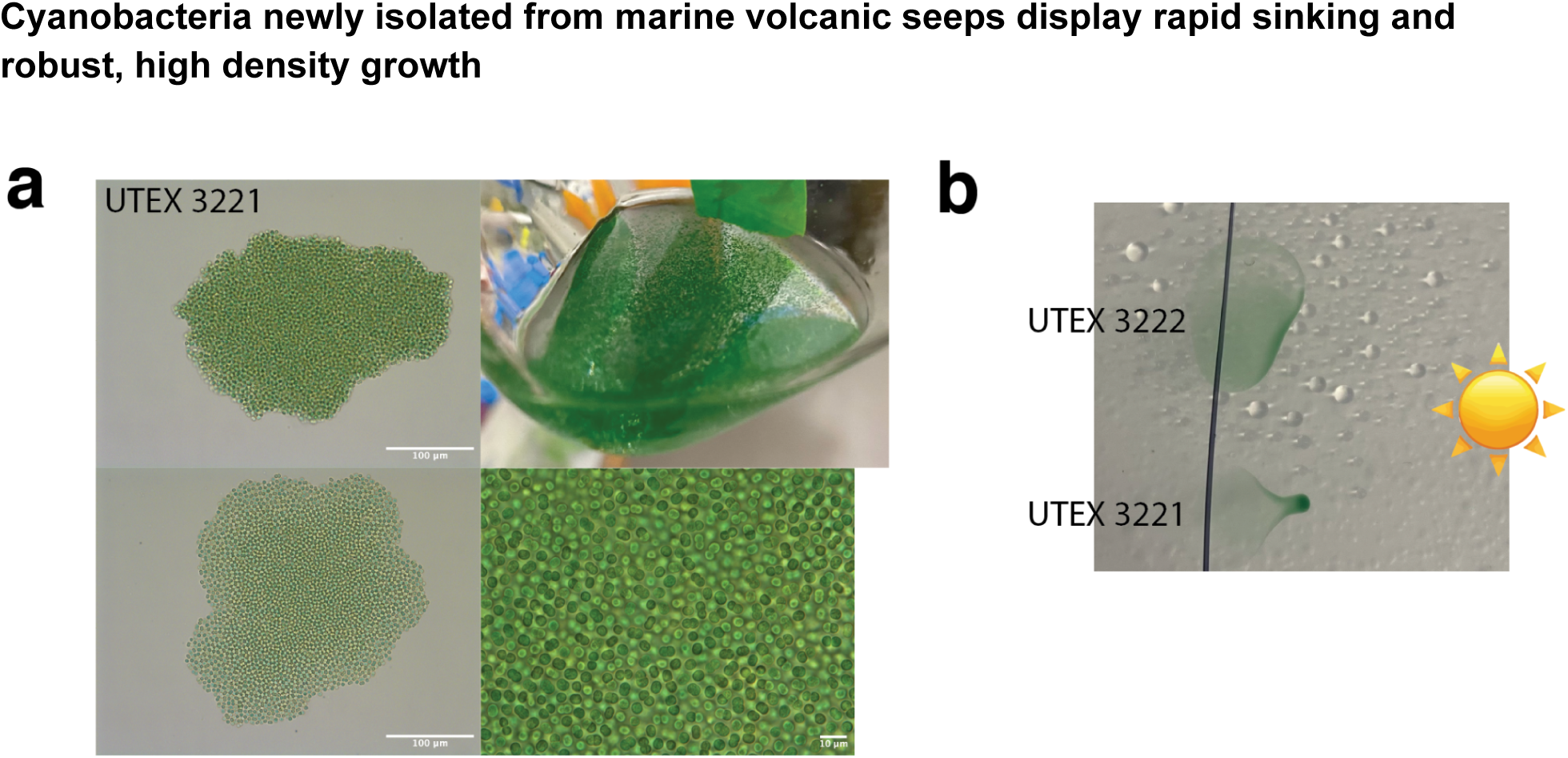
a) Micrograph of UTEX3222 b) Macroscopic aggregates visible when growing UTEX3221 in liquid medium c) Micrographs showing further detail of the aggregates in (b). d) Micrograph showing UTEX3221 aggregate with higher magnification.

**Supplemental Figure 2.**
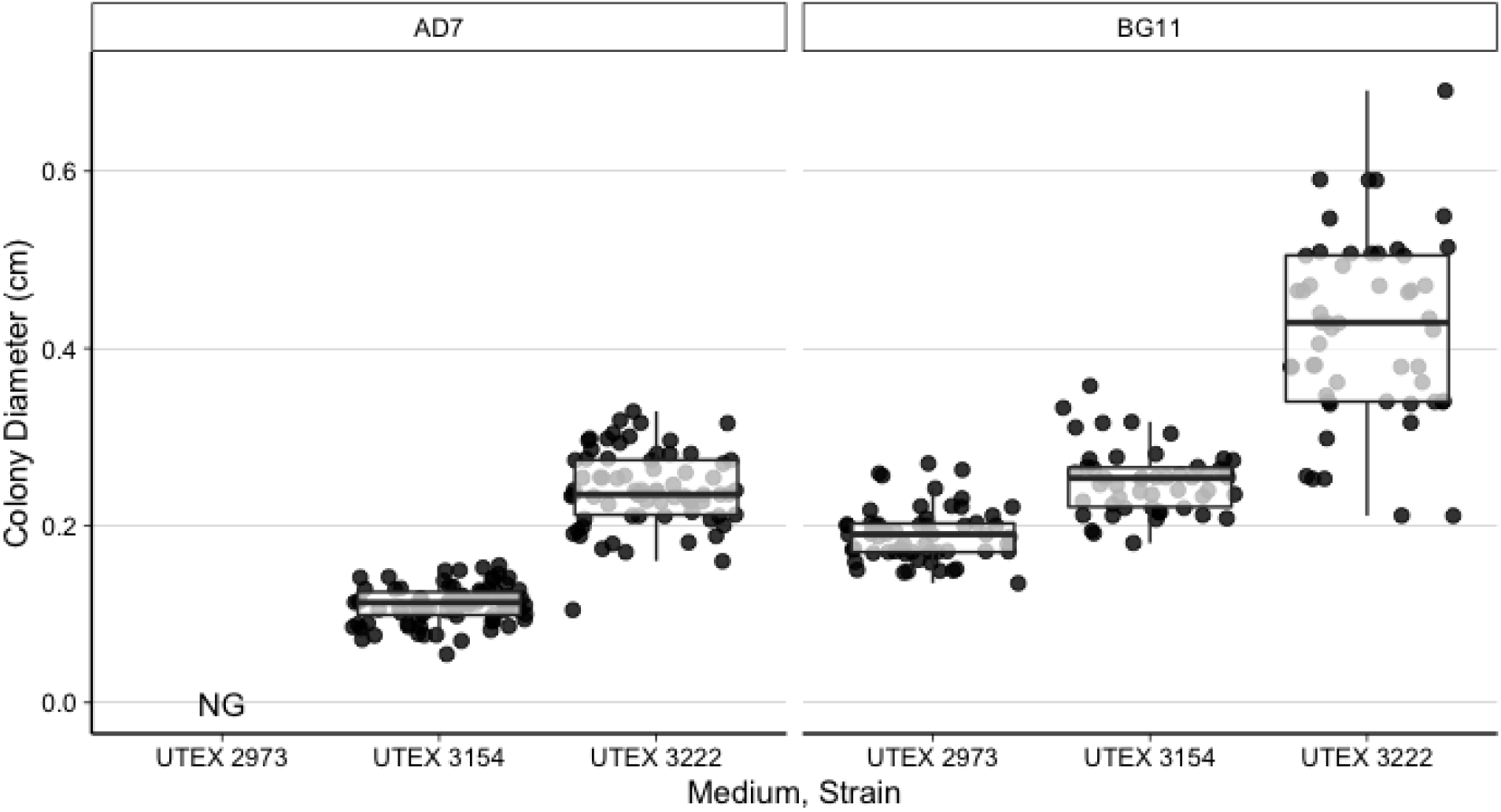
Comparison of colony sizes observed in Figure 1B. Measurements of individual colonies are depicted by points, and additionally summarized with box plots in the style of Tukey. No Growth (NG) was observed for UTEX 2973 on AD7 medium. All media additionally supplemented with 4mg/L vitamin B-12 in this experiment.

**Supplemental Figure 3.**
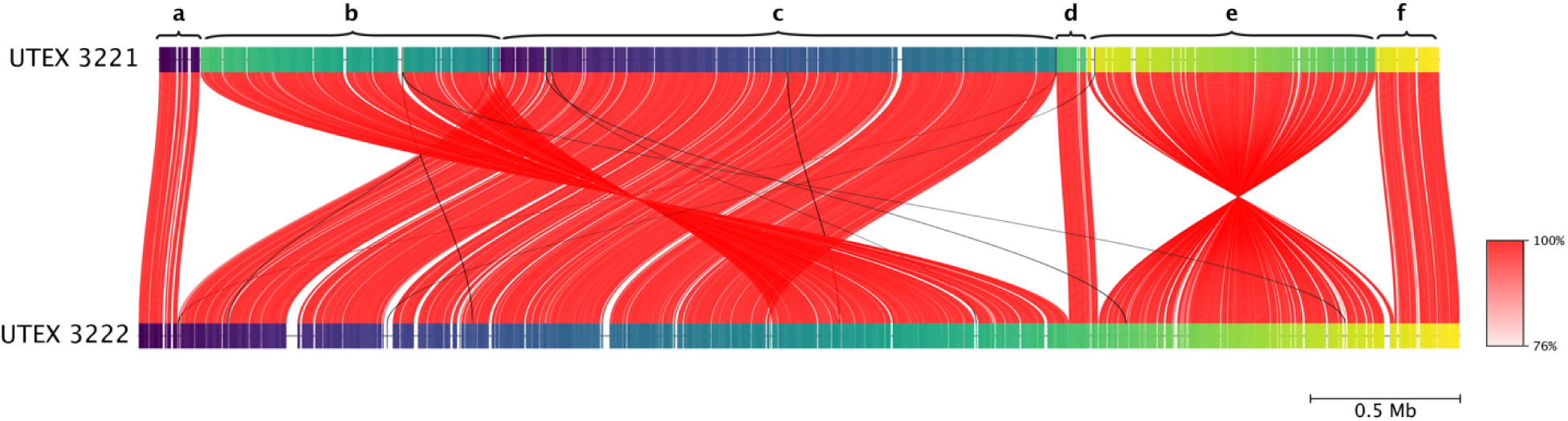
A visualization of alignment between UTEX 3222 and UTEX 3221 genomes. Red lines indicate reciprocal mapping segments, and are colored by Average Nucleotide Identity. One interpretation is that the strains differ by translocation and inversion of segment b, as well as inversion of segment e.

**Supplemental Figure 4.**
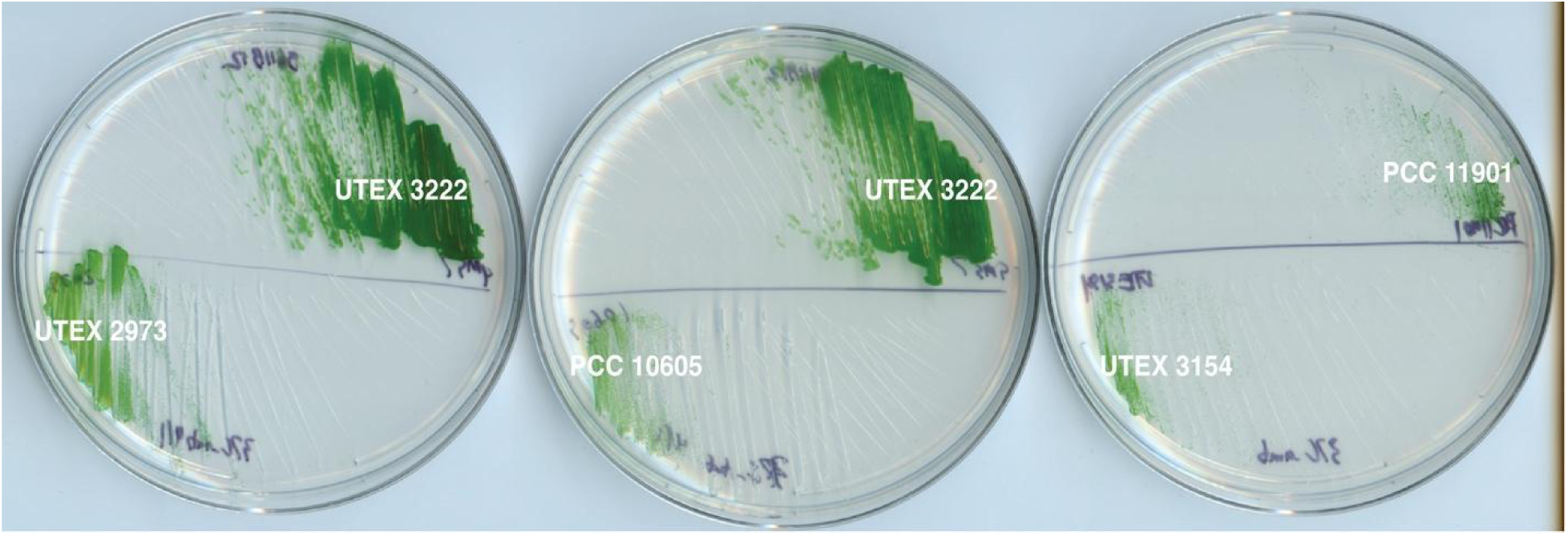
A panel of cyanobacteria strains grown on BG11+B12 medium in ambient air conditions, 37C, approximately 100uE light, for 4 days.

**Supplemental Figure 5:**
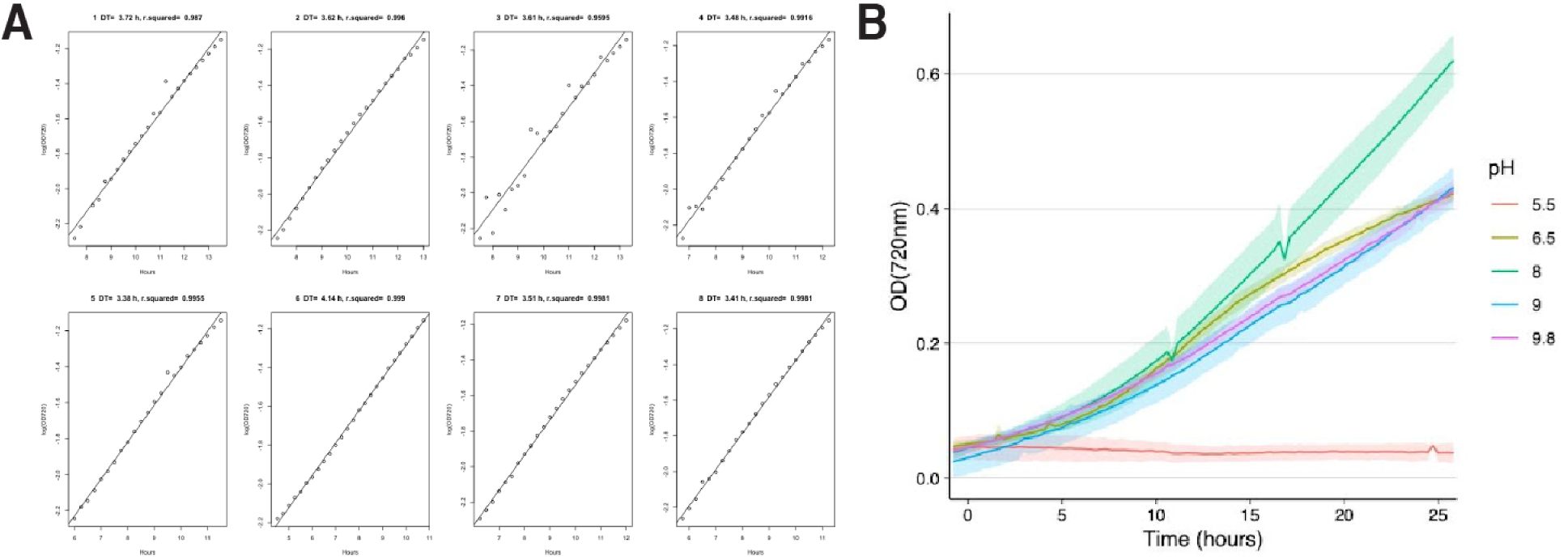
Additional information on exponential growth A) Representative linear regressions used to derive exponential growth rate (µ). Individual optical density (OD) measurements are plotted on a logarithmic axis against time. Linear regression is depicted as a line, and doubling times and r^2^ values of fits are given. B) Growth curves in varying pH medium. Lines depict the mean value across replicates, and shaded areas depict the standard deviation across replicates.

**Supplemental Figure 6:**
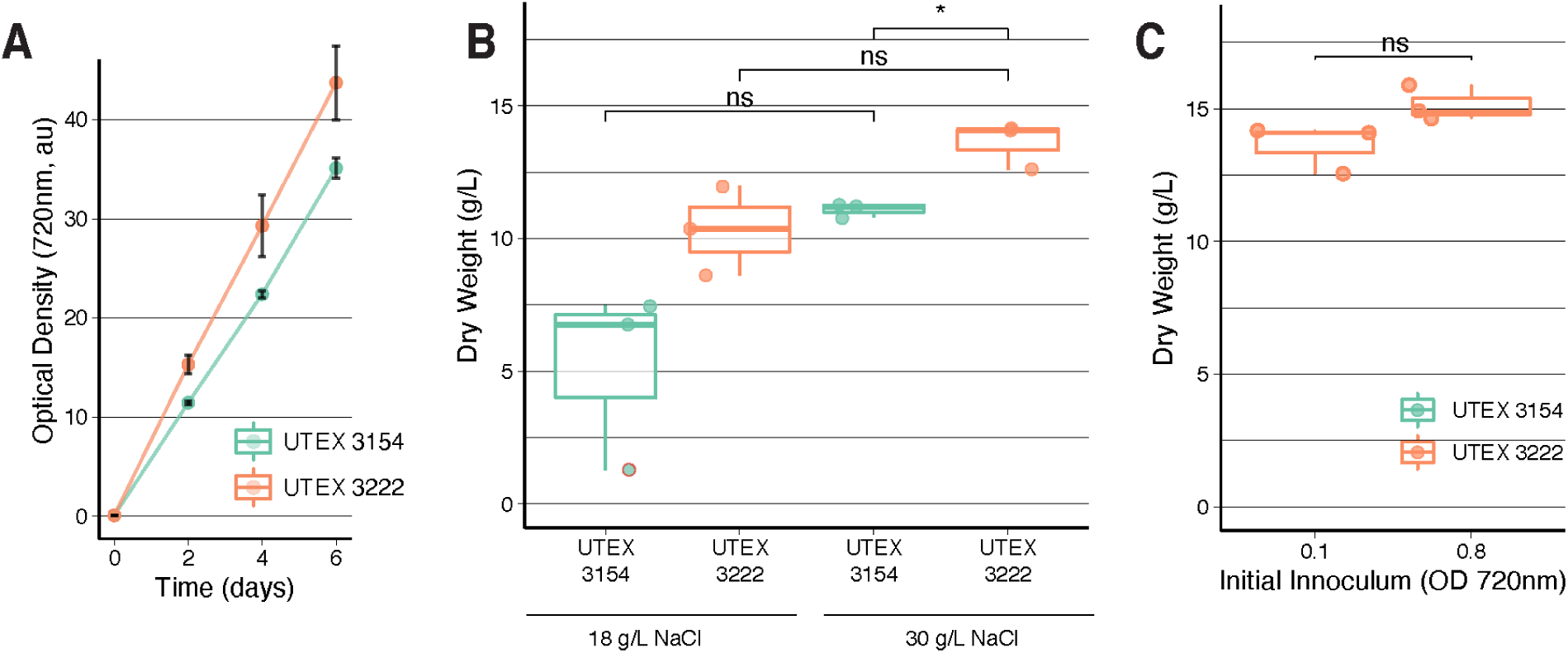
Additional High Density Growth A) Optical Density (OD720) monitored over the course of high density batch growth, 0.5% CO2, 200µE light. Points depict the mean of triplicate measurements, with error bars depicting the Standard Error. B) Comparison of 18g/L NaCl medium to 30g/L NaCl medium. Conditions were 0.5% CO_2_ and 200µE light, 37C, 7 days. C) Comparison of inoculation density. Conditions were as above, except with 5% CO2 supplied. Triplicate experiments are depicted by points, and summarized by boxplots. Data in A 18g/L NaCl are reproduced from Figure 3B. Unpaired t-tests were performed, yielding either non-significant (ns) or p <0.05 (*) results depicted. P-values reading left to right are 0.093, 0.057, and 0.031 for B, and 0.078 for C.

**Supplemental figure 7:**
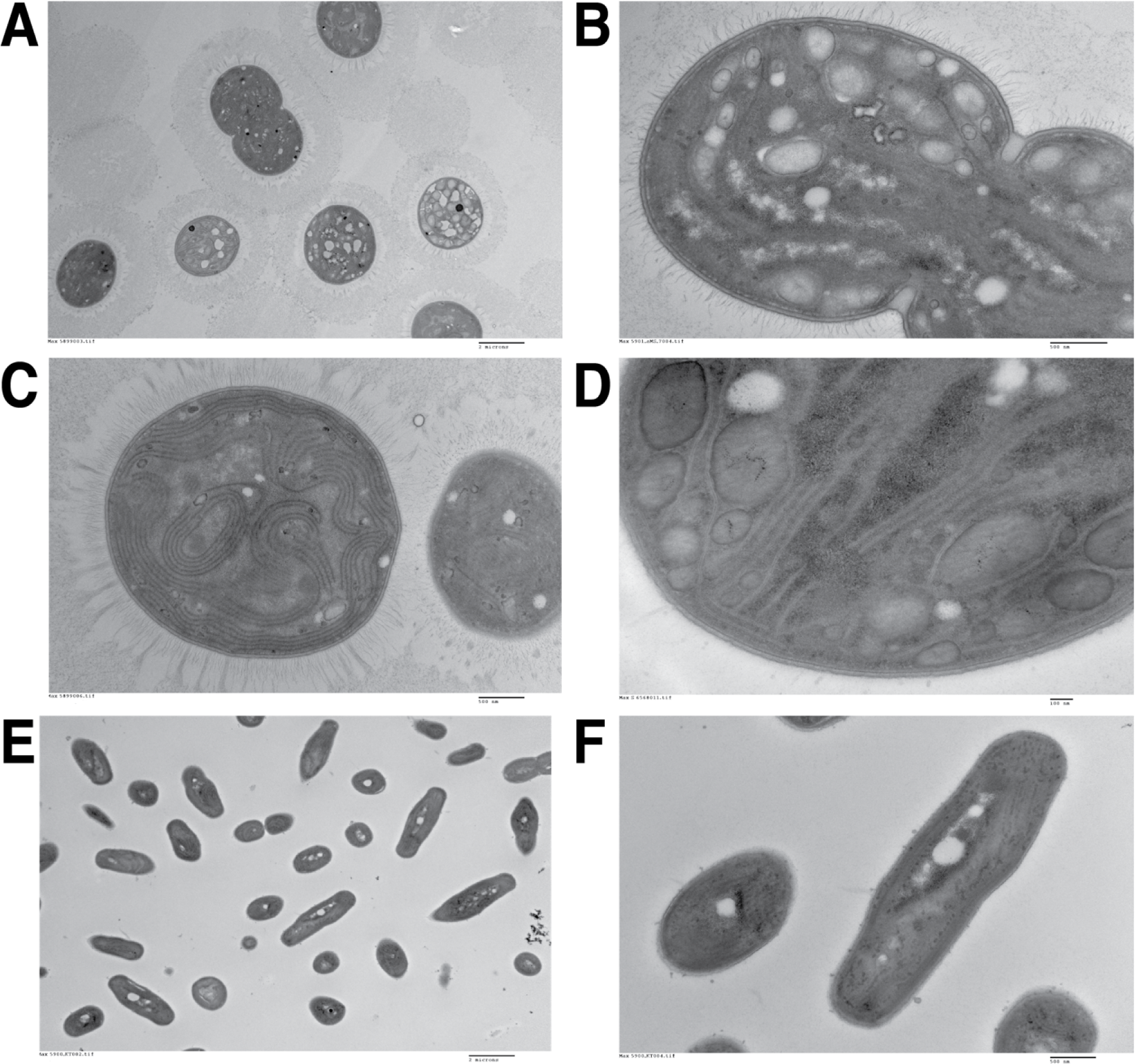
Additional featured Transmission Electron Micrographs A) UTEX 3222 at low magnification, showing extracellular material, and a subset of cells featuring putative storage granules B) UTEX 3222, Higher magnification of a dividing cell featuring putative storage granules C) UTEX 3222 with thylakoid membranes and putative pili visible. D) UTEX 3222, higher magnification of a cell with both putative storage granules and thylakoid membranes visible. E) UTEX 3154 for comparison, displaying smaller cells and relative lack of visibly staining extracellular material.

**Supplementary Figure 8:**
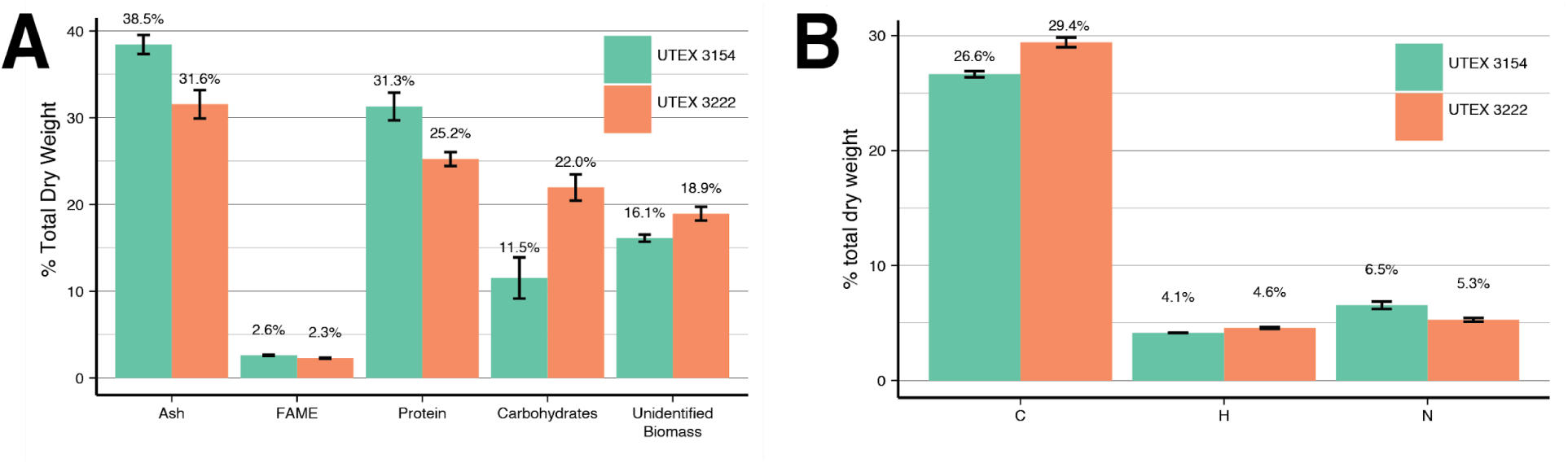
further summary of biomass characterization. A) Composition of major macromolecules as a percentage of Total Dry Weight, contrasting with Ash-Free Dry Weight (AFDW) in Figure 4B. B) C/H/N Elemental composition of UTEX 3154 and UTEX3222 biomass. Bars depict the mean of three replicate experiments, and error bars depict the standard error of these measurements.

**Supplemental Figure 9:**
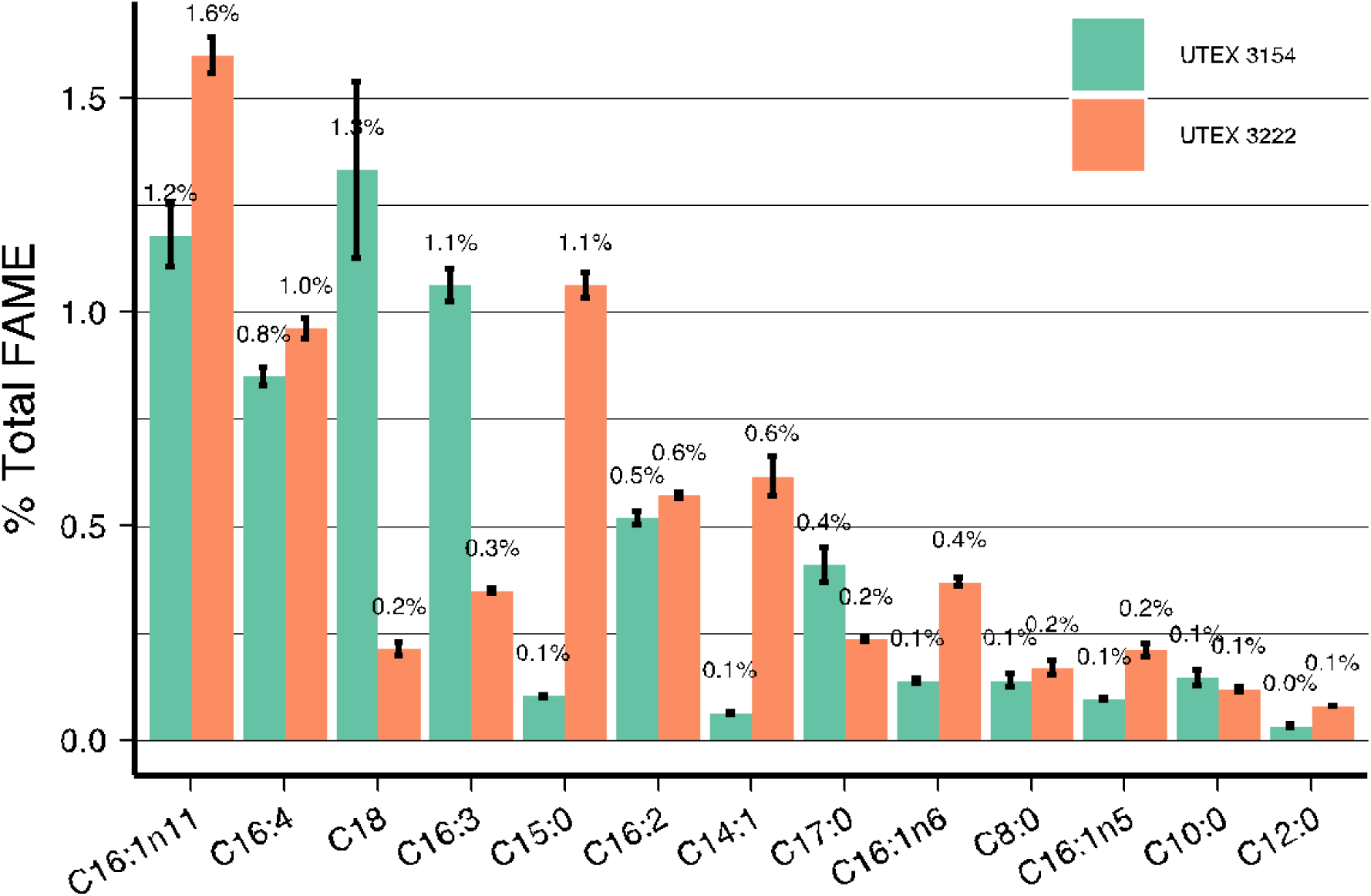
comparison of FAME abundance for FAME species

**Supplemental Figure 12:**
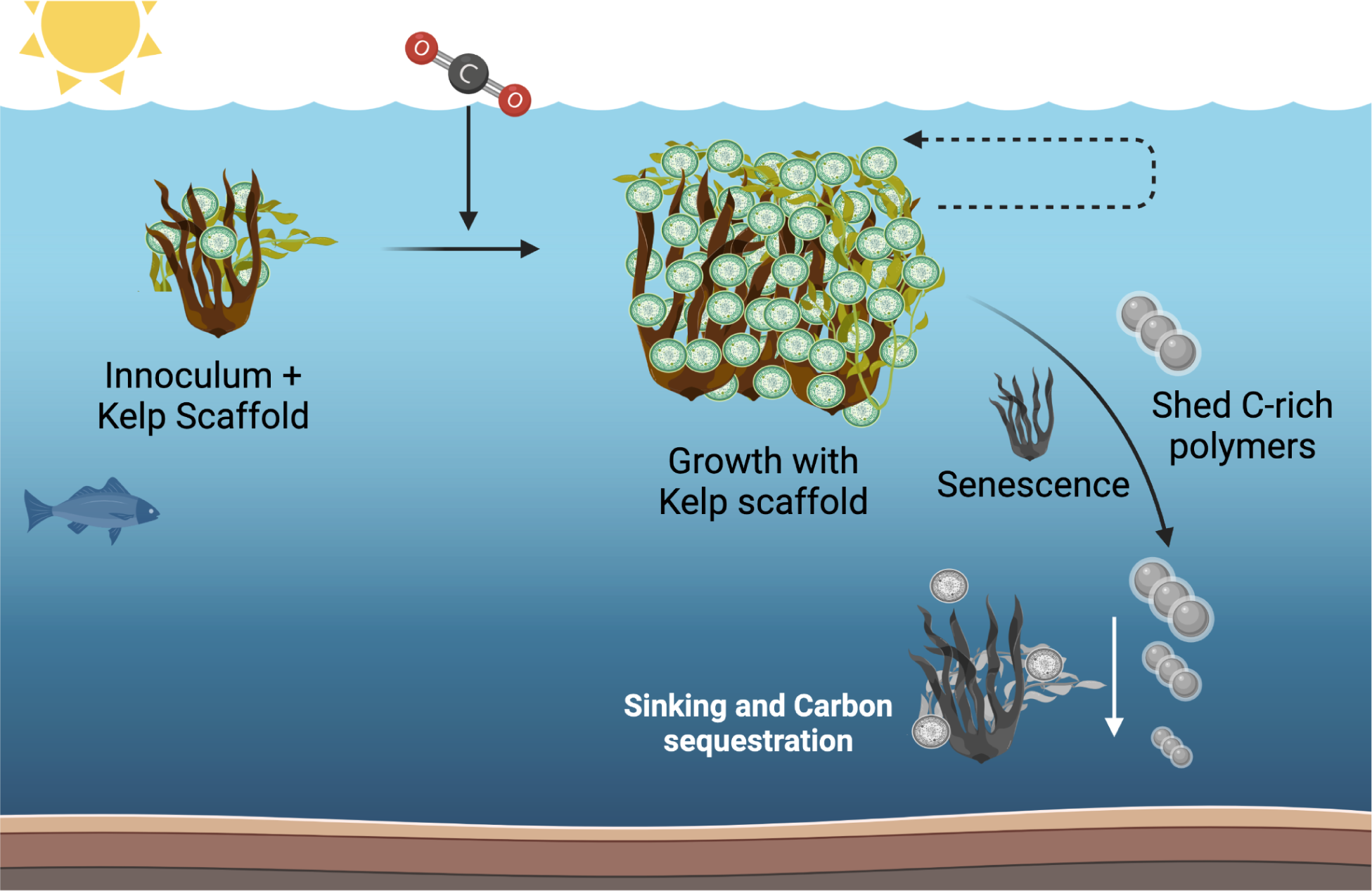
A schematic of a potential marine carbon-sequestration system using cyanobacteria and a kelp scaffold. Kelp potentially offer a floating scaffold on which cyanobacteria can grow and photosynthesize in the photic zone for extended time periods before sinking. The cyanobacteria characterized in this study produce abundant EPS, which may allow them to adhere to kelp. Potentially this system could be self-sustaining and expand over time. Ideally, such a system would produce a substantial quantity of C-rich polymers, by shedding EPS or other exudates. This would maximize the carbon leaving the system, while potentially minimizing Nitrogen, Phosphorous, or other nutrients being co-sequestered alongside Carbon. Such polymers would ideally be long-lived in the marine environment. It’s possible that the lifetime of such systems may be finite-there may be a process of senescence and sinking for both kelp scaffold and cyanobacteria (greyscale).

